# Molecular determinants of Arc oligomerization and formation of virus-like capsids

**DOI:** 10.1101/667956

**Authors:** Maria Steene Eriksen, Oleksii Nikolaienko, Erik Ingmar Hallin, Sverre Grødem, Helene J. Bustad, Marte Innselset Flydal, Rory O’Connell, Tomohisa Hosokawa, Daniela Lascu, Shreeram Akerkar, Jorge Cuéllar, James J. Chambers, Ian Merski, Gopinath Muruganandam, Remy Loris, Tambudzai Kanhema, Yasunori Hayashi, Margaret M. Stratton, José M. Valpuesta, Petri Kursula, Aurora Martinez, Clive R. Bramham

## Abstract

Expression of activity-regulated cytoskeleton-associated protein (Arc) is critical for long-term synaptic plasticity, memory formation, and cognitive flexibility. The ability of Arc to self-associate and form virus-like capsid structures implies functionally distinct oligomeric states. However, the molecular mechanism of Arc oligomerization is unknown. Here, we identified a 28-amino-acid region necessary and sufficient for Arc oligomerization. This oligomerization region is located within the second coil of a predicted anti-parallel coiled-coil in the N-terminal domain (NTD). Using alanine scanning mutagenesis, we found a 7-amino-acid motif critical for oligomerization and Arc-mediated transferrin endocytosis in HEK cells. Intermolecular fluorescence lifetime imaging in hippocampal neurons confirmed self-association mediated by the motif. To quantify oligomeric size, we performed a single-molecule photobleaching analysis of purified Arc wild-type and mutant. This analysis revealed a critical role for the NTD motif in the formation of higher-order Arc oligomers (30-170 molecules). Moreover, assembly of higher-order wild-type Arc oligomers was significantly enhanced by addition of GFP RNA. Purified wild-type Arc formed virus-like capsids, as visualized by negative-stain EM, and was estimated by light scattering analysis to contain 40-55 Arc units. In contrast, mutant Arc formed a homogenous dimer population as demonstrated by single-molecule TIRF imaging, size-exclusion chromatography with multi-angle light scattering analysis, small-angle X-ray scattering analysis, and single-particle 3D EM reconstruction. Thus, the dimer appears to be the basic building block for assembly. Herein, we show that the NTD motif is essential for higher-order Arc oligomerization, assembly of virus-like capsid particles, and facilitation of oligomerization by exogenous RNA.

**SIGNIFICANCE:** Arc protein is rapidly expressed in neurons in response to synaptic activity and plays critical roles in synaptic plasticity, postnatal cortical developmental, and memory. Arc has diverse molecular functions, which may be related to distinct oligomeric states of the protein. Arc has homology to retroviral Gag protein and self-assembles into retrovirus-like capsid structures that are capable of intercellular transfer of RNA. Here, we identified a motif in the N-terminal coiled-coil domain of mammalian Arc that mediates higher-order oligomerization and formation of virus-like capsids. The basic building block is the Arc dimer and exogenous RNA facilitates further assembly. The identified molecular determinants of Arc oligomerization will help to elucidate the functional modalities of Arc in the mammalian brain.

## INTRODUCTION

Activity-regulated cytoskeletal associated protein (Arc) has emerged as a key regulator of neuronal plasticity, memory, and postnatal cortical development (Bramham et al., 2010; Nikolaienko et al., 2018; Shepherd and Bear, 2011). Arc is induced as an immediate early gene, and the neuronal activity-induced Arc RNA and protein are subject to rapid turnover, indicating a highly dynamic mode of action. Loss of function studies support a role for Arc in long-term potentiation (LTP) and long-term depression (LTD) of synaptic transmission and homeostatic synaptic scaling. These diverse responses are mediated by distinct Arc protein-protein interaction complexes in the postsynaptic dendritic compartment and the neuronal nucleus (Chowdhury et al., 2006; DaSilva et al., 2016; Korb et al., 2013; Nair et al., 2017; Okuno et al., 2012; Zhang et al., 2015). Thus, convergent lines of evidence support a role for Arc as a signaling hub protein and cell-autonomous organizer of synaptic plasticity (Nikolaienko et al., 2018).

The mechanisms that dictate Arc function at the molecular level are poorly understood. Post-translational modification of Arc by SUMOylation (Craig et al., 2012; Nair et al., 2017) and ERK-catalyzed phosphorylation (Nikolaienko et al., 2017) are implicated in the regulation of protein-protein interactions and subcellular localization, while Arc turnover is regulated by ubiquitination, acetylation, as well as GSK-catalyzed phosphorylation (Gozdz et al., 2017; Greer et al., 2010; Lalonde et al., 2017; Mabb et al., 2014; Wall et al., 2018). In addition, biochemical studies show that recombinant Arc protein is capable of reversible self-association (Byers et al., 2015; Myrum et al., 2015). It has also been shown that purified human Arc forms higher-order oligomeric species, dependent on ionic strength, but reverses to monomers and dimers at low ionic strength (Myrum et al., 2015). This reversible oligomerization, as opposed to nonspecific aggregation, raised the possibility that Arc function is related to its oligomeric state.

Recent advances highlight a structural and functional relationship between Arc and retroviral Gag polyprotein. Arc was identified in a computational search for domesticated retrotransposons harboring Gag-like protein domains (Campillos et al., 2006). Biochemical studies showed that mammalian Arc has a positively charged N-terminal domain (NTD) and a negatively charged C-terminal domain (CTD), separated by a flexible linker (Myrum et al., 2015). Crystal structure analysis of the isolated CTD revealed two lobes, both with striking 3D homology to the capsid (CA) domain of HIV Gag (Zhang et al., 2015). In retroviruses, self-association of CA allows assembly of Gag polyproteins into the immature capsid shell (Lingappa et al., 2014; Perilla and Gronenborn, 2016). Remarkably, recombinant Arc from fruit fly and rat was subsequently shown to self-assemble into spheroid particles with resemblance to HIV Gag capsids (Ashley et al., 2018; Pastuzyn et al., 2018). The Arc capsids are released in extracellular vesicles and capable of transmitting RNA cargo to recipient cells (Ashley et al., 2018; Pastuzyn et al., 2018). These discoveries implicate Arc as an endogenous neuronal retrovirus, and oligomeric assembly of Arc into virus-like capsids mediates the capture and intercellular transfer of RNA (Parrish and Tomonaga, 2018; Shepherd, 2018).

A recent structural analysis of full-length monomeric human Arc shows a compact shape, in which the oppositely charged NTD and CTD interact (Hallin et al., 2018). Interestingly, Drosophila Arc has a CA-like CTD domain but lacks the NTD found in mammals (Zhang et al., 2015). The Arc NTD is expected to have evolved from the matrix (MA) domain of the Gag polyprotein (Campillos et al., 2006). There is also a functional similarity, as both the Arc NTD and the Gag MA domain mediate binding to phospholipid membranes (Hallin et al., 2018; Mailler et al., 2016). However, important differences may exist, as structural analysis of the Arc NTD indicates an antiparallel coiled-coil, which is not present in retroviral MA (Hallin et al., 2018).

A major goal is to elucidate the relationship between Arc as a dynamic mediator of intracellular signaling *versus* its newly described function as a virus-like capsid. We therefore sought to identify molecular determinants of Arc self-association and assembly into capsids. Our approach combines biochemical and physicochemical methods (affinity purification, *in situ* protein crosslinking, size-exclusion chromatography (SEC), dynamic light scattering (DLS)), as well as fluorescence lifetime imaging, single-molecule photobleaching analysis, and electron microscopy (EM). We identified a 28-amino-acid stretch in the Arc NTD that is both necessary and sufficient for oligomerization. The critical oligomerization region is located within the second coil of the predicted anti-parallel coiled-coil. Alanine scanning mutagenesis identified a 7-amino-acid motif underlying oligomerization above the dimer stage and leading to capsid formation, as validated by EM. Surprisingly, similarly to the nucleocapsid domain of HIV Gag, Arc oligomerization is greatly enhanced by exogenous RNA in a manner dependent on the second coil motif. Our findings shed light on the molecular mechanisms of mammalian Arc oligomerization and capsid assembly.

## RESULTS

### Arc oligomerization is mediated by the NTD second coil

First, we sought to map the regions of the Arc protein mediating oligomerization using a stringent affinity purification assay. We co-expressed two variants of Arc in a human embryonic kidney cell line (HEK293FT):1) Arc with an N-terminal fusion to mTurquoise2 (mTq2) and 2) Arc with a a C-terminal fusion to StrepII tag (Fig.1A). Next, mTq2-fused full-length Arc (1-396) was coprecipitated with StrepII-tagged Arc, indicating complex formation (Fig. 1C). Specificity was confirmed by the absence of mTq2-positive bands in purifications from cells transfected with StrepII-Arc and the empty mTq2 vector. Arc contains five cysteine residues (C34, C94, C96, C98, C159), which we suspected might form disulfide-linked oligomeric species. However, substituting all five cysteines with serines did not affect the Arc-Arc interaction, showing that oligomerization is not a result of disulfide bond formation (Fig. 1C). We next co-expressed truncated versions of Arc fused to mTq2 to determine the minimal regions needed for association (Fig.1B). When StrepII-tagged full-length Arc was coexpressed with mTq2-fused N-terminal region (1-140), linker (135-216) or C-terminal region (208-396), only the N-terminal region showed binding (Fig. 1D). The involvement of the N-terminal region in self-association agrees with our previous SAXS data showing that the isolated Arc CTD is fully monomeric (Hallin et al., 2018).

**Fig. 1.**
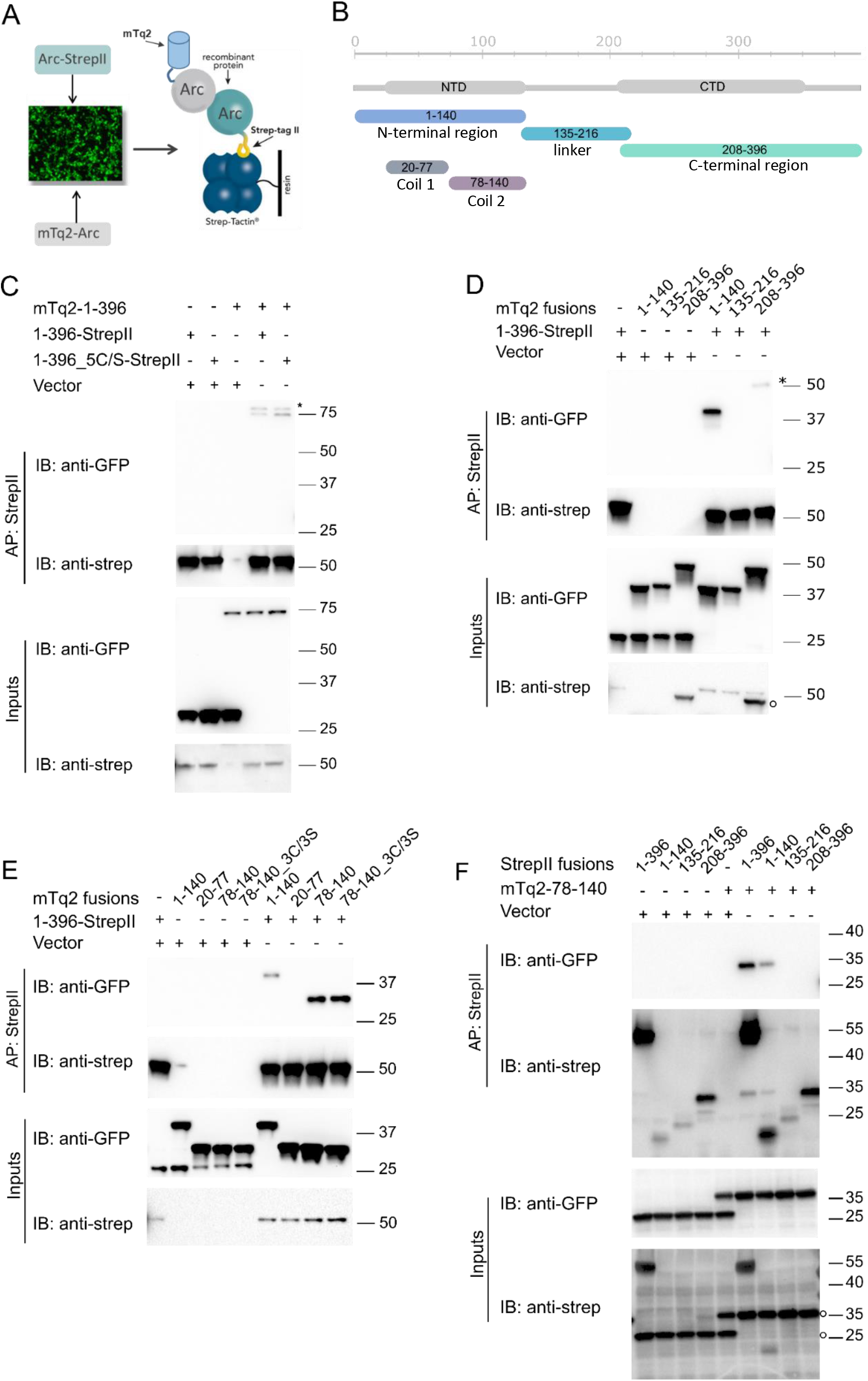
Arc self-oligomerization is mediated by the N-terminal domain second coil. (A) Model showing the principle behind Arc affinity purification. (B) Schematic representation of Arc fragments used in the study. (C) mTurquoise2-fused (mTq2) full-length Arc was coexpressed in HEK293FT cells with StrepII-tagged full-length Arc^WT^ or a five-site cysteine mutant. mTq2 (vector) was used as control. (D) StrepII-tagged full-length Arc was coexpressed with mTq2-fused Arc fragments (mTq2 fusions) as indicated by the numbers: 1-140, 135-216 and 208-396. (E) StrepII-tagged full-length Arc was coexpressed with mTq2-fused Arc fragments (mTq2 fusions; 1-140, 20-77, 78-140 and a cysteine to serine mutant of 78-140). (F) mTq2-fused Arc 78-140 was coexpressed with StrepII-tagged Arc as indicated by residue numbers (StrepII fusions; 1-140, 135-216, 208-396). (C-F) Cell lysates were incubated with Strep-Tactin Sepharose and the bound proteins eluted in sample buffer. Proteins were detected after SDS-PAGE and Western blot analysis by anti-GFP and anti-Strep antibodies. Asterisks indicate unspecific bands with molecular weight too high compared to inputs. Open circles indicate remaining anti-GFP immunofluorescence from first round of membrane probing.

According to secondary structure predictions, the Arc NTD has a predicted coiled-coil region which can be divided into two alpha coils, with residues 20-77 and 78-140 corresponding to the first and second coil, respectively. Our previous structural work indicates an elongated, anti-parallel coiled-coil structure of the NTD (Hallin et al., 2018). We therefore examined co-purification of full-length Arc with each of the NTD coils and found that only the second coil (residues 78-140) interacted with Arc (Fig. 1E). Similar binding was obtained with a Cys to Ser mutant (3C/3S, residues 94, 96, and 98) of the second coil, again ruling out formation of disulfide-linked oligomers (Fig 1E). The second coil co-purified with the isolated Arc NTD but not with the isolated linker region or CTD (Fig. 1F). Furthermore, high-affinity binding between StrepII-tagged 78-140 and mTq2-tagged 78-140 was observed, demonstrating self-association of the NTD second coil (*SI Appendix, Fig. S1*).

### Identification of an oligomerization motif in the Arc NTD second coil

Next, we sought to delineate which residues within the NTD second coil (residues 78-140) mediate Arc-Arc interaction. We first generated a series of deletion mutants of the second coil in which 18 amino acid stretches were removed, starting with every 9^th^ consecutive residue, giving an overlap of 9 residues between the deleted regions (*SI Appendix, Table S1*). Deletion mutants Δ78-95 and Δ114-131 did not bind full-length Arc (Fig. 2A), however, the Δ78-95 mutant was associated with decreased expression of the peptide. We then generated substitution variants, in which non-overlapping stretches of seven amino acids were mutated to alanine (Fig. 2B, *SI Appendix, Table S1*). With the exception of s78-84A, which exhibited impaired expression, the substitution variants expressed at levels similar to wild-type (WT) control. Affinity purification analysis showed that all substitutions within the central region of the second coil, from residue Gln99 to Trp126, inhibited binding of the second coil to full-length Arc (Fig. 2B). Importantly, oligomerization was similarly inhibited or abolished when mutations were introduced into full-length Arc (Fig.2C). Most notably, alanine substitution of the sequence _113_MHVWREV_119_ (Arc^s113-119A^) abolished binding to Arc-StrepII.

**Fig. 2.**
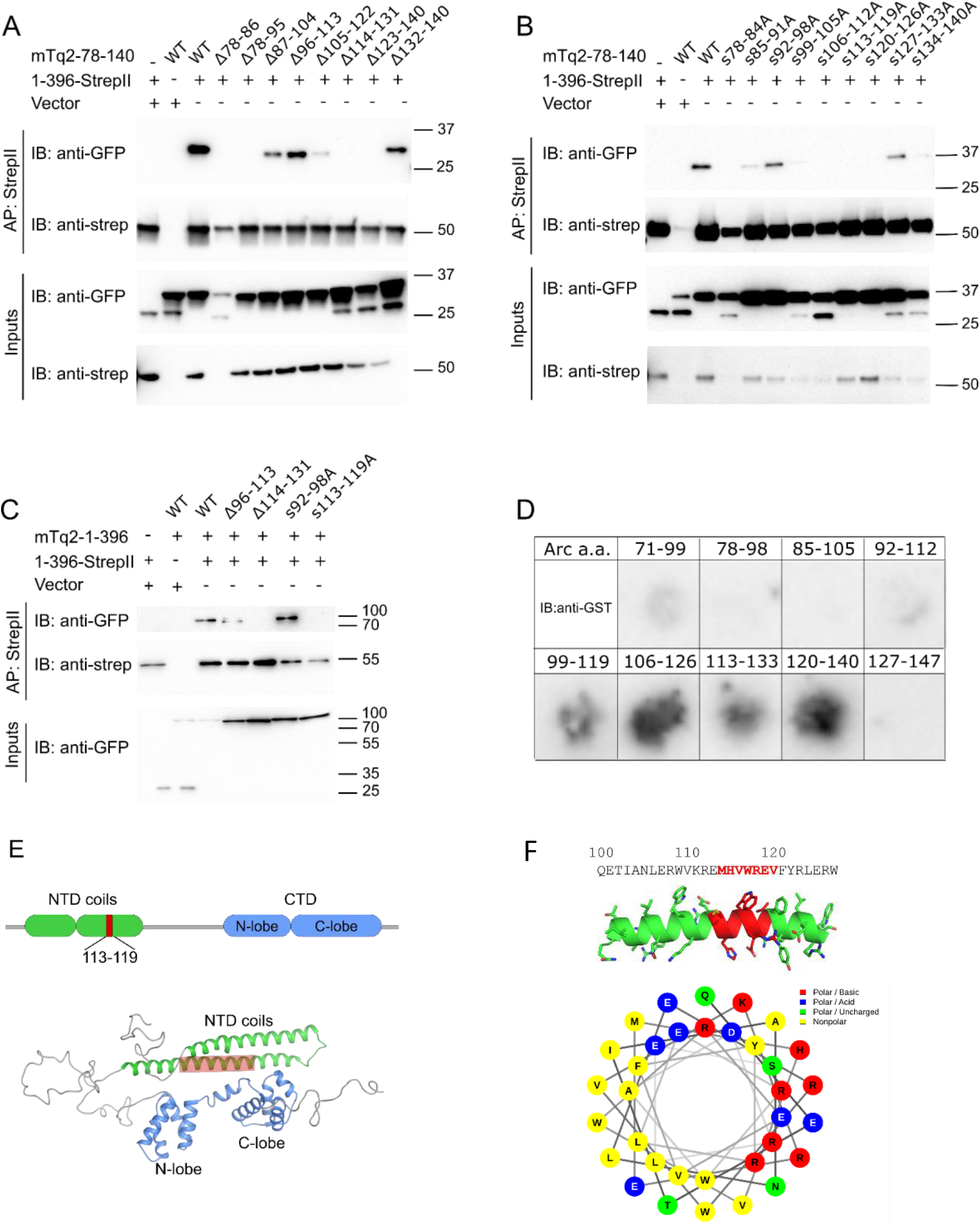
Identification of an oligomerization interface in the Arc NTD second coil. (A) StrepII-tagged full-length Arc was coexpressed with mTq2-fused Arc 78-140 wild-type or Arc 78-140 with 18 amino acids deleted as the numbers indicate. (B) StrepII-tagged full-length Arc was coexpressed with mTq2-fused Arc 78-140 wild-type or Arc 78-140 with 7-amino acid alanine substitutions as indicated. (C) StrepII-tagged full-length Arc was coexpressed with mTq2-fused full-length Arc with either deletions or substitutions as indicated. In panel A-C, cell lysates were incubated with Strep-Tactin Sepharose and the bound proteins eluted in sample buffer. Proteins were detected after SDS-PAGE and Western blot analysis by anti-GFP and anti-Strep antibodies. (D) A peptide array consisting of 21-mer Arc peptides was incubated with GST-fused Arc 78-140. Probing with anti-GST revealed binding of Arc 78-140 to peptides spanning amino acids 99-126. E, top) Cartoon shows position of the oligomerization region and the 7-amino acid motif in the NTD second coil. (E, bottom) Oligomerization region in full-length Arc 3D structure (SAXS hybrid model (Hallin et al., 2018). (F, top) Oligomer peptide sequence with 113-119 motif in red. (F, bottom) Helical wheel representation of the amino acids in oligomerization region shows the amphipathic nature of the peptide. The amino acid charge distribution in full-length Arc is shown in SI Appendix, Fig. S2.

As a complementary approach, we mapped the NTD second coil interaction site using a peptide-tiling array of Arc region 71-147. Each spot on the array displays a 21-residue peptide, with 7-residue overlap between sequences. The array was incubated with purified GST-fused Arc 78-140 (complete second coil) and immunoblotted with anti-GST antibodies. Binding was observed with tiling peptides spanning Arc region 99-126 (Fig. 2D). Pure GST did not bind to immobilized peptides. These *in vitro* binding results identify an oligomeric interface in the second coil and corroborate findings from affinity purification of tagged Arc from HEK cells.

Taken together, Arc region 99-126 is necessary and sufficient for Arc oligomerization (Fig. 2E and 2F). We refer to this 28-amino-acid stretch as the oligomerization region, while the critical Arc 113-119 sequence is referred to as the oligomerization motif. The position of the oligomerization region in the context of the SAXS hybrid 3D model of the Arc monomer (Hallin et al. 2018) is shown in Fig 2E. The oligomerization region is amphipathic with hydrophobic and hydrophilic residues on opposite sides of the predicted helix (Fig. 2F). Notably, the oligomerization region has an isoelectric point of 8.6 and is flanked by patches of dense positive charge in the highly basic NTD (pI of 9.6; *SI Appendix, Fig. S2*).

### The Arc oligomerization region forms hexamers

The oligomerization region is predicted to adopt an α-helical conformation and self-associate. To assess secondary structure and detect possible oligomeric forms of the oligomerization region, we performed circular dichroism (CD) spectrum analysis, SAXS, and SEC with multiangle light-scattering (SEC-MALS) analysis on a synthetic peptide (residues 99-132) containing the defined oligomerization region (99-126) and six additional C-terminal residues, which based on our analysis of substitution mutants may contribute to oligomerization. The CD spectrum indicated high α-helical content as expected (*SI Appendix, Fig. S3A*). SEC showed the formation of low-order oligomers by the peak position at an elution volume of ~16 ml (*SI Appendix, Fig. S3B*) and a lack of larger complexes. MALS indicated a size of 27 kDa, corresponding to a hexamer. SAXS analysis of the same peptide also showed a hexameric assembly and an elongated shape with P32 symmetry (*SI Appendix, Fig. S3C-F)*. These experiments demonstrate a strong intrinsic property of the peptide to oligomerize in the absence of cellular factors, such as protein binding partners, membranes, or nucleic acids.

### Fluorescence lifetime FRET imaging in hippocampal slices confirms self-association of the second coil mediated by the oligomerization motif

To assess the second coil-mediated interaction in live neurons, we performed fluorescence lifetime FRET imaging (FLIM-FRET) in CA1 pyramidal cells of organotypic rat hippocampal slice cultures. Slices were transfected by gene gun with plasmids expressing the second coil (residues 78-140) fused to GFP (donor) and mCherry (acceptor) (Fig. 3A). In FLIM-FRET, a decrease in fluorescence lifetime of the GFP donor, as measured by time-correlated single-photon counting, indicates increased protein-protein interaction. The fluorescence lifetime of GFP-Arc second coil WT was equal to the positive control, a GFP-mCherry fusion that gives constitutive FRET (Fig. 3B), but significantly lower than the negative control (GFP-P2A-mCherry) in which the two fluorophores are separated by a self-cleaving peptide. Next, we examined the role of the 113-119 oligomerization motif in mediating the interaction between coils. When the mutant coil was used as the FRET acceptor, the interaction was significantly reduced compared to WT control (Fig. 3B). These live-imaging data show strong self-association of the second coil in hippocampal neurons and confirm the critical role of the 113-119 motif in oligomerization. However, we also note that the FRET signal in the second coil mutant did not reach the level of the negative control, indicating residual oligomerization.

**Fig. 3.**
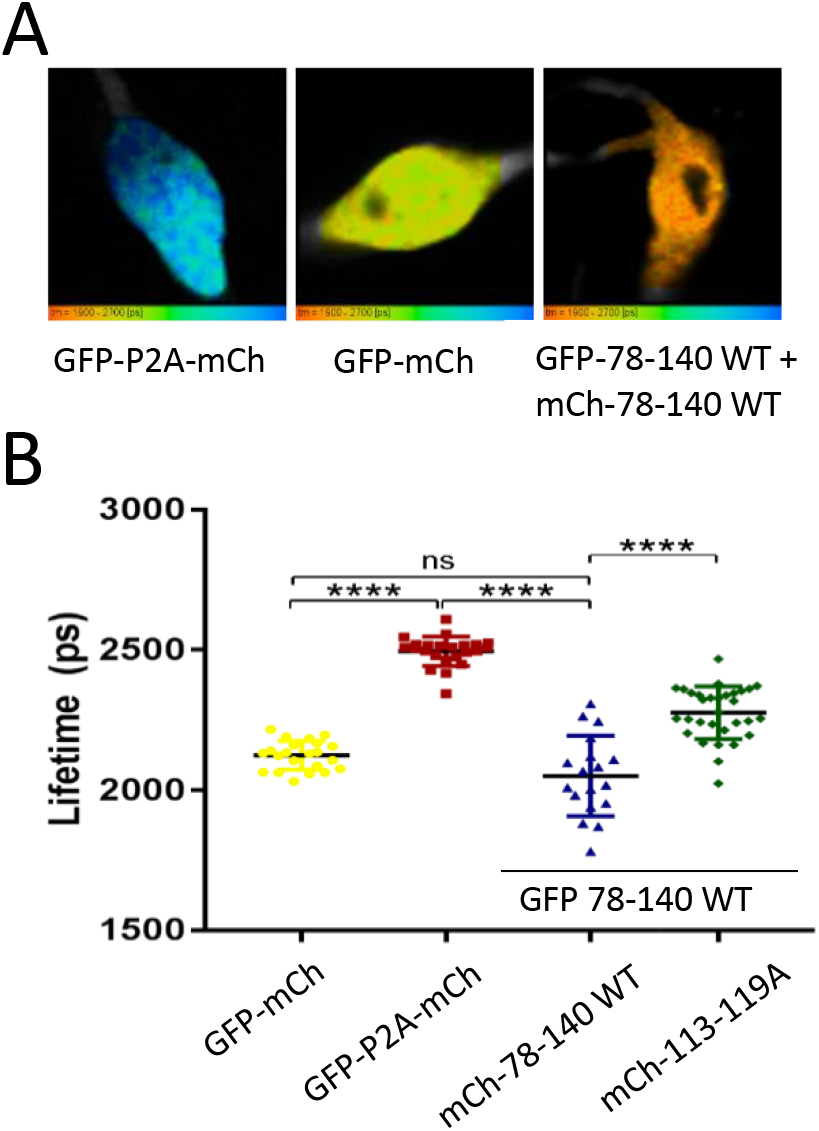
Fluorescence lifetime FRET imaging of Arc second coil oligomerization in CA1 pyramidal cells of organotypic rat hippocampal slice cultures. (A) Representative FLIM-FRET images of neurons expressing negative control (GFP-P2A-mCh), positive control (GFP-mCh) and Arc 78-140 (GFP-78-140_WT + mCh-78-140_WT). (B) Lifetime distribution of analyzed cells. Lines indicate mean and standard deviations. One-way ANOVA between groups (P<0.05) with Tukey’s multiple comparisons post-hoc correction. n = 22, 23, 18, 28, 31, respectively.

### *In situ* protein crosslinking reveals persistence of Arc dimer in oligomerization motif mutant

The findings from hippocampal slices suggested that alanine substitution of the 113-119 motif does not completely abolish Arc interactions, which contrasts with our findings from affinity purification of Arc expressed in HEK cells. We therefore considered that cell lysis and homogenization might disrupt weak oligomeric interactions, as previously shown for α-synuclein (Dettmer et al., 2013). To detect oligomeric species *in situ*, we examined Arc interactions in live HEK cells exposed to protein crosslinkers. Cells were transfected with constructs expressing mTq2-fused Arc^WT^, Arc^s92-98A^, Arc^s113-119A^, or empty vector. In the main protocol used, cells were treated for 10 min with 100 μM disuccinimidyl glutarate (DSG) before harvesting (Fig. 4). A strong Arc (and GFP) immunoreactive band was detected at 75 kDa, corresponding to monomeric mTq2-tagged Arc (Fig. 4A). In crosslinked preparations only, Arc immunoreactive bands were detected at approximately 180 kDa and 280 kDa, corresponding to dimers and trimers, respectively (Fig 4A). The dimer was prominent for all constructs, and there was no difference between constructs in the dimer/monomer ratio, indicating no impact of the 113-119 motif at the level of dimer formation (Fig. 4B). However, the trimer/dimer ratio was significantly reduced in Arc^s113-119A^ relative to both Arc^s92-98A^ and Arc^WT^ (Fig. 4B). These results reveal prominent cellular expression of an Arc dimer and further imply a specific role for the Arc 113-119 motif in the assembly of oligomers above the dimer stage.

**Fig 4.**
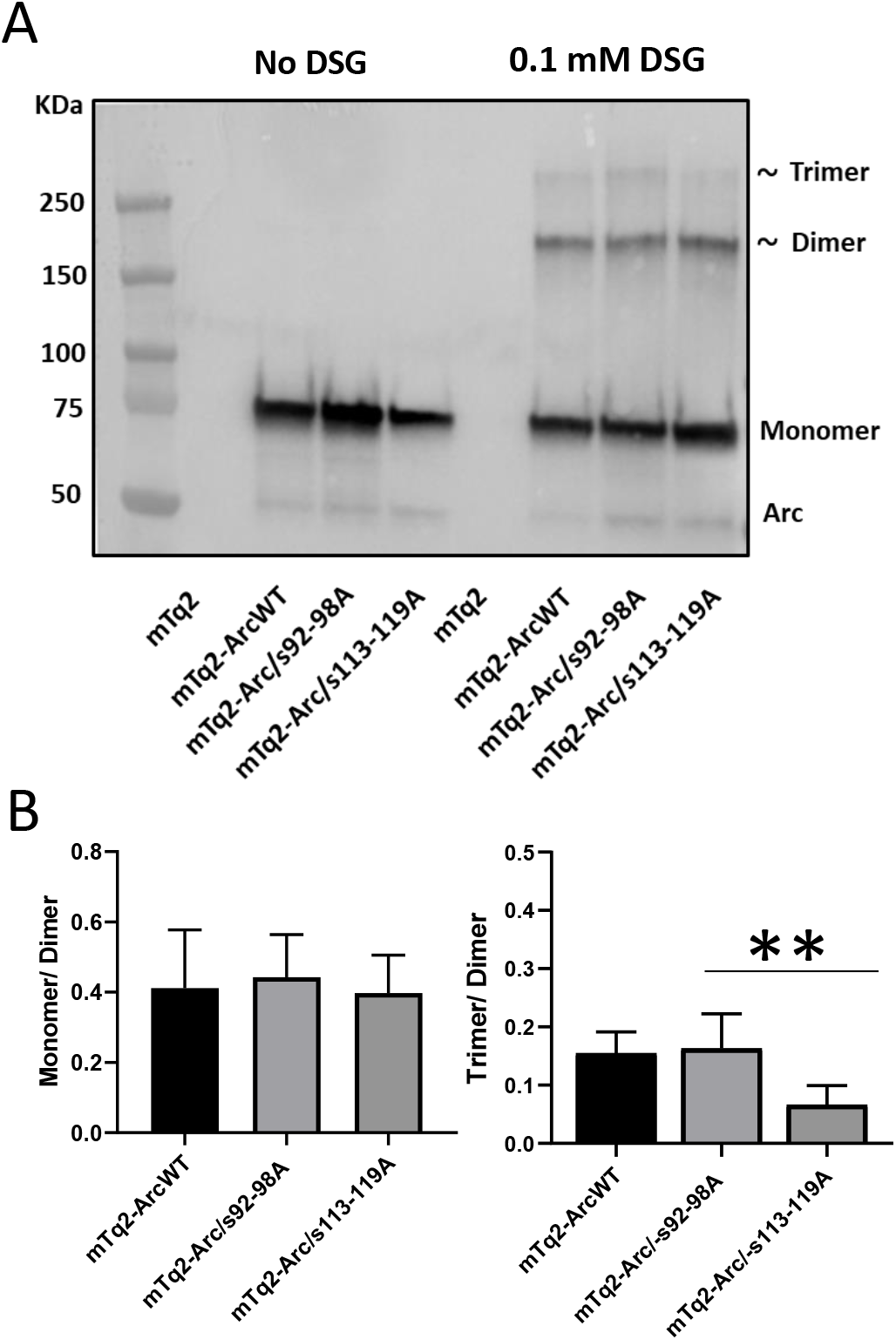
In situ protein crosslinking reveals prominent Arc dimer but inhibition of trimer formation in Arc 113-119 second coil mutant. HEK293FT cells were transfected with Arc^WT^, Arc^s92-98A^ or Arc^s113-119A^, all tagged at the N-terminus with mTurquoise2. For crosslinking, cells were treated with disuccinimidyl glutarate (DSG) dissolved in DMSO to a final concentration of 0.1 mM for 10 min. (A) Arc immunoreactive bands were detected at approximately 75 kDa, 180 kDa, and 280 kDa corresponding to the monomer, dimers, and trimers of mTq2-tagged Arc. Oligomers were detected only in crosslinked samples. Similar results were obtained with paraformaldehyde crosslinking (not shown). (B) Bar graphs show ratio of Arc monomer/dimer expression (left) and trimer/dimer expression (right). Values and mean + S.E.M. One-way ANOVA with Bonferroni multiple comparison was used for statistical analysis **P<0.01.

### Mutation of Arc oligomerization domain impairs transferrin endocytosis

Arc is an adaptor protein for clathrin-mediated endocytosis and facilitates cellular uptake of transferrin (Chowdhury et al., 2006). The Arc NTD second coil harbors an endophilin binding site determinant (residues 89-100) involved in endocytosis (Chowdhury et al., 2006), and the same region contains a cysteine cluster _94_CLCRC_98_ involved in palmitoylation and membrane targeting of Arc (Barylko et al., 2017). We therefore explored whether oligomerization mediated by the NTD second coil contributes to endocytosis, as assessed by uptake of Alexa Fluor 647-conjugated transferrin in HEK cells. As expected, expression of Arc^WT^ enhanced transferrin uptake relative to empty vector control (*SI Appendix, Fig. S4*). Expression of the endophilin binding site mutant, Arc^s92-98A^, failed to increase transferrin uptake (*SI Appendix, Fig. S4*), although the protein successfully forms oligomers, as shown by affinity purification (Fig 2B and 2C). Notably, the oligomerization-deficient Arc mutants (Δ114-131 and s113-119A) failed to facilitate transferrin uptake (*SI Appendix, Fig. S4*). As levels of monomer and dimer expression did not differ between Arc^s113-119A^, Arc^s92-98A^ and Arc^WT^, these results support a role for oligomerization above the dimer stage in promoting Arc endocytic activity, which also requires the endophilin-binding site.

### Single-molecule imaging of Arc oligomeric state: critical role of NTD second coil in RNA-induced oligomerization

In order to rigorously quantify the oligomeric state of Arc, we employed single-molecule total internal reflection fluorescence (smTIRF) microscopy. For these experiments, we used recombinantly purified rat Arc with an N-terminal SNAP tag and a C-terminal AviTag. Purified Arc was labeled with an AlexaFluor488 SNAP substrate, C-terminally biotinylated and affixed onto glass cover slides through streptavidin binding (Fig. 5A). The size of each Arc oligomer was determined by photobleaching, where the number of photobleaching steps corresponds to the number of dye molecules, and therefore the number of Arc monomers in each oligomer (Fig. 5B).

**Fig. 5.**
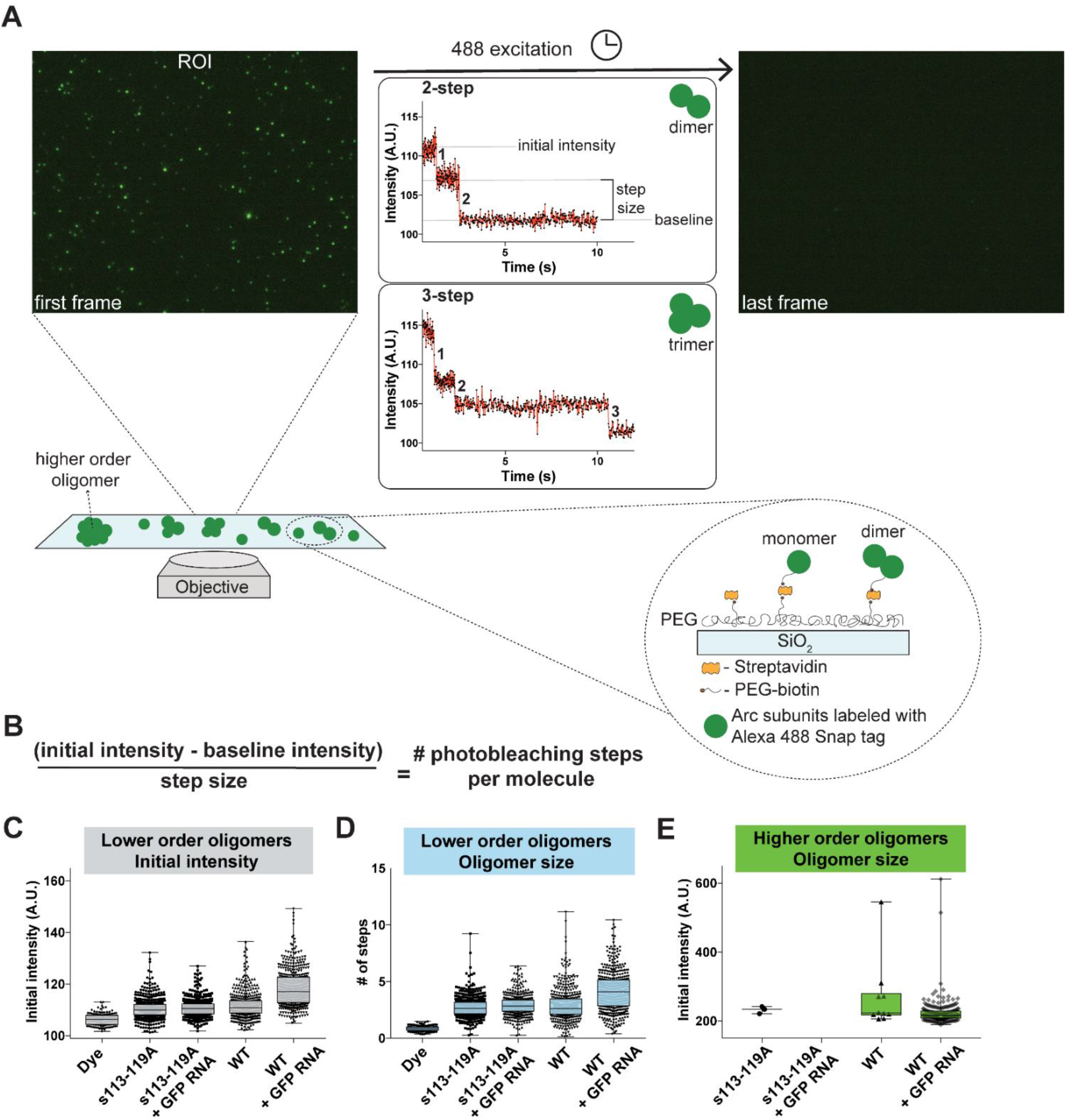
Single-molecule analysis of Arc reveals critical role of second coil motif in RNA-induced higher-order oligomerization. (A) Flowchart of the photobleaching experiment using single-molecule TIRF microscopy. Biotinylated Arc is affixed to glass cover slides using a PEG-biotin/streptavidin linkage, as shown. A region of interest (ROI) is exposed to constant 488 nm excitation, causing molecules to photobleach over time. Examples of the first and last frames from a photobleaching video are shown. If the single molecule is an Arc dimer, 2 discrete photobleaching steps are observed. Examples of 2-step and 3-step photobleaching are shown. (B) Equation used to calculate number of photobleaching steps per molecule. (C) We compared the initial intensity (first frame of ROI) of single molecules of several samples and conditions (see X-axis label). Molecules with an initial intensity less than ~160 A. U. were considered lower-order oligomers. Box and whisker plots show the 10^th^−90^th^ percentile data points for clarity. (D) After calculating the step size from each experiment (see A), the initial intensity was converted to number of steps (see B). The number of photobleaching steps for the lower-order oligomers is plotted. The number of steps correlates to the size of oligomer (*i.e.*, 4 photobleaching steps indicates a tetrameric complex). (E) For the same samples as in B, we separately analyzed molecules brighter than 160 A. U., which were considered higher-order oligomers. The initial intensity of all molecules that qualified is plotted.

Arc samples were imaged using smTIRF, where each fluorescent spot corresponds to one oligomeric complex (Fig. 5A). Spots with clearly identified steps of greater than 2 arbitrary units (A. U.) over 5 frames were used to calculate the step size for each experiment (see examples in Fig. 5A). The number of photobleaching steps per spot was then determined by dividing the total intensity by the calculated step size (Fig. 5B). The initial intensity is proportional to the number of fluorophores per spot (Fig. 5C). From these initial intensities, we calculated the number of photobleaching steps, which suggest that lower-order Arc oligomers of fewer than 10 molecules exist predominantly as dimeric and trimeric species. In Tris buffer (pH 7.5), Arc^WT^ had a median step size of 2.61 (±1.42 S.D., 0.068 S.E.M.). Similarly, the Arc^s113-119A^ mutant also showed the presence of lower-order oligomeric species with an average step size of 2.64 (± 0.89 S.D., 0.038 S.E.M.). These data show that Arc is mainly dimeric/trimeric in Tris buffer.

Arc capsid formation was previously shown to be facilitated by addition of EGFP mRNA (Pastuzyn et al., 2018), but the mechanism is unknown. Interestingly, studies of Gag lattice assembly show that non-specific binding to RNA promotes Gag multimerization (Lingappa et al., 2014; Muriaux et al., 2002). We speculated that exogenous RNA might similarly facilitate Arc oligomerization. To address this question, exogenous EGFP mRNA was added to purified Arc protein before attaching it to coverslips. When EGFP mRNA was added to purified Arc^WT^, the median step size increased to 4.1 (±1.77 S.D., 0.084 S.E.M.). When EGFP mRNA was added to Arc^s113-119A^ mutant, there was no shift in oligomeric size observed. Thus, lower-order Arc oligomerization is facilitated by exogenous RNA and this response depends on the Arc 113-119 second coil motif.

We separately analyzed discrete brighter fluorescent spots, which correspond to higher-order oligomeric species. It is not reliable to determine the number of steps in the higher-order oligomers due to the large number of clustered fluorophores. This results in limited discrete steps, which confounds our analysis. However, if we assume that the step size is the same, given that the fluorophore is the same, we estimate the size of the higher-order oligomers is between 30-170 monomers of Arc per complex. Arc aggregation on the slides was sometimes detected, and these very bright amorphic regions were not analyzed. The majority (84%) of the higher-order oligomers fall in the range of ~40-70 subunits per complex. In the absence of EGFP mRNA, very few higher-order oligomers were found in the Arc^WT^ and Arc^s113-119A^ samples (10 and 3 oligomers, respectively) (Fig 5E). However, when EGFP mRNA was added to Arc^WT^, there was a 19-fold increase in the number of higher-order oligomers to 190 oligomers. In striking contrast, zero higher-order oligomers were observed in the Arc^s113-119A^ samples with mRNA added. The findings show that higher-order oligomerization of Arc is greatly enhanced by exogenous GFP RNA through a mechanism that requires the 113-119 oligomerization motif.

### The NTD second coil motif mediates formation of Arc capsids

A key question is whether the higher-order oligomers driven by the Arc NTD assemble into capsids. In our first analysis of recombinant human Arc by negative-stain EM, we found irregular particles of ~30 nm diameter in samples that were prepared in Hepes buffer with high ionic strength (Myrum et al., 2015). Here, using PBS buffer in sample preparation, we show that wild-type human Arc forms rounded, viral-like capsids (Fig. 6) similar in size and shape to those reported for recombinant rat and Drosophila Arc (Ashley et al., 2018; Pastuzyn et al., 2018). The majority of capsids have a diameter of ~30 nm, although there is heterogeneity in size. In agreement with Pastuzyn et al. (Pastuzyn et al., 2018), addition of EGFP mRNA to purified protein resulted in a more regular, smoother capsid perimeter (Fig. 6A-D).

**Fig. 6.**
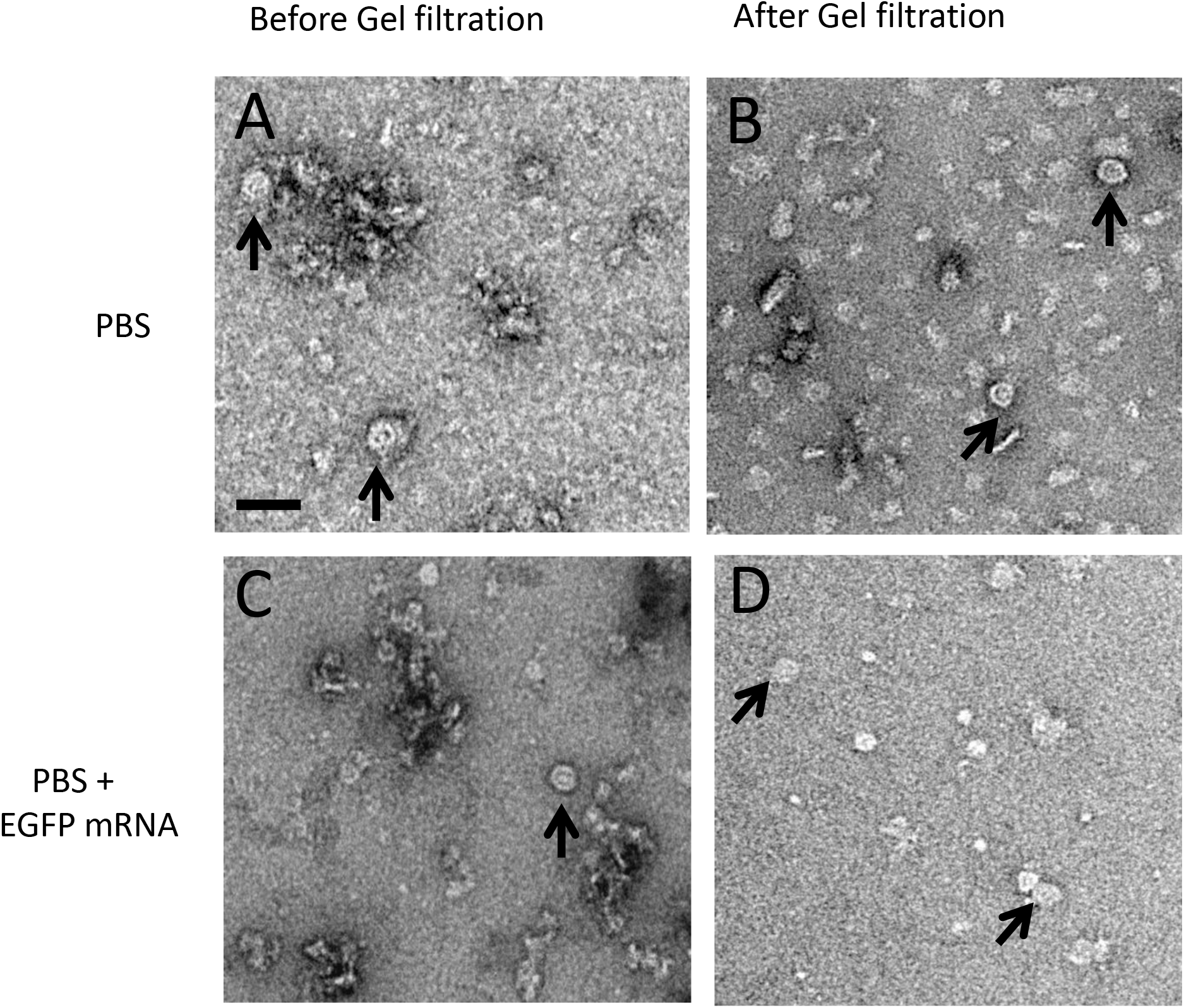
Negative-stain electron microscopy and the effect of GFP RNA. Transmission electron microscopy images of negatively stained Arc preparations: (A) in PBS buffer, without mRNA; (B) in PBS buffer, without mRNA, after SEC; (C) with mRNA before SEC. (D) with mRNA after SEC. The four samples show a clear heterogeneity, with particles of different sizes, aggregates, but also spherical capsid-like structures of ~30 nm diameter (arrows). In most cases, capsids contain an electron-dense substance. Scale bar, 60 nm.

Next, we used DLS to estimate the size of purified Arc^WT^ and Arc^s113-119A^. Both proteins display one population by volume, but with 40-45% polydispersity (Fig. 7A). However, the Z-average values for Arc^WT^ of 28.1±11.9 nm includes particles with the capsid diameter observed by EM and corresponds to an estimated molecular mass between 900 and 2400 kDa (1.6±0.7 MDa). Based on these DLS data we propose a unit number between of 40 and 55 Arc molecules per capsid. In contrast, Arc^s113-119A^ did not display a population of this size, but had a mean diameter of 11.4±4.7 nm, corresponding to an estimated molecular mass between 100 and 300 kDa (Fig. 7A). The mutant eluted as a single peak on SEC, and MALS confirmed a mass of 106 kDa, corresponding to a homodimer (Fig. 7B).

**Fig. 7.**
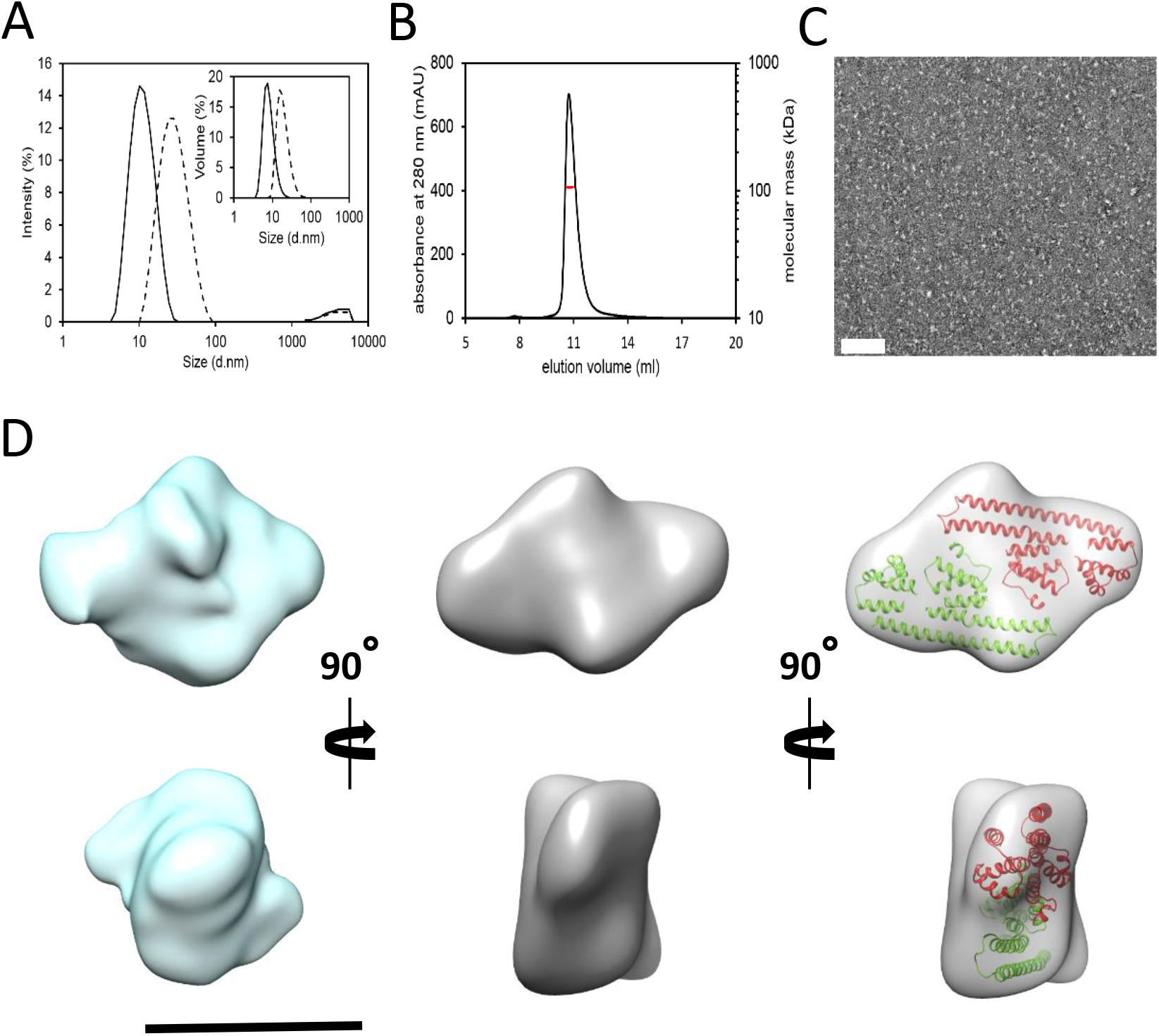
The Arc second coil motif mediates formation of virus-like capsids. (A) DLS analysis with size-estimation of purified Arc^WT^ (dashed line) and Arc^s113-119A^ (solid line) based on intensity and volume (inset) size distribution. (B) SEC-MALS analysis of the Arc^s113-119A^ mutant. (C) Electron microscopy image of negative-stained Arc^s113-119A^ mutant (right). Bar indicates 50 nm. (D) Two orthogonal views of the 3D reconstruction of the Arc^s113-119A^ mutant (left) without symmetry imposition, (center) with C2 symmetry imposed and (right) the latter one with the docking of two copies (red and green) of the atomic structures of the N-terminal coiled-coil and the two CTD lobes, the N-terminal (4×3i.pdb) and the C-terminal one (4×3x.pdb) (Hallin et al., 2018). The docking shows that two Arc monomers can fit in the reconstructed volume, but does not necessarily represent the actual structure of the dimer.

Importantly, negative-stain EM analysis of the mutant revealed a homogeneous population of small elongated particles (Fig. 7C), with no sign of capsid-like structures observed for Arc^WT^ (Fig. 6B). We performed single-particle 3D EM reconstruction of 55,057 automatically selected particles from the Arc^s113-119A^ sample. The structure obtained, with no symmetry imposition, has a rhomboid, symmetrical shape (Fig. 7D, left). The size (~130×85 Å) and the apparent symmetry suggest that the particles are formed by a dimer of the Arc^s113-119A^ mutant. A subsequent refinement with C2 symmetry rendered a similar volume (Fig. 7D, center), and docking of the SAXS-based atomic structure of monomeric Arc (Hallin et al. 2018) fits with a dimer (Fig. 7D, right).

SAXS analysis of the mutant independently gave a dimeric structure closely resembling the 3D EM reconstruction, with a slightly more elongated shape (*SI Appendix, Fig. S5*). An elongated dimer shape was also obtained with Arc^s113-119A^ with an N-terminal MBP fusion tag, and the external positions of MBP in the SAXS structure indicate an outward orientation of the NTD in the dimer (*SI Appendix, Fig. S5*). We conclude that capsid formation is mediated by the NTD second coil, and mutation of the Arc 113-119 motif blocks assembly above the dimer stage.

## DISCUSSION

Arc is recognized as an enigmatic, master regulator of synaptic plasticity and memory of importance for human cognition. Arc is both a signaling hub and capable of forming retrovirus-like capsids for transmission of RNA in extracellular vesicles. The present work elucidates molecular determinants of Arc oligomerization and formation of capsid-like structures. Convergent evidence from biochemical and biophysical approaches, single-molecule imaging, and EM reveals a critical role for the Arc NTD coiled-coil in oligomerization. We identified an oligomeric interface of 28 amino acids located within the second coil of the predicted anti-parallel coiled-coil of the Arc NTD. Alanine scanning further identified a 7-amino-acid motif, 113MHVWREV119, critical for higher-order oligomerization above the dimer stage and leading to capsid formation. Higher-order self-association was strongly facilitated by exogenous RNA and dependent on the Arc^113-119^ motif.

The ancient retroviral origins of Arc invites a comparison of the structural and functional properties of the Arc NTD with the extensively studied domains of retroviral Gag. The Gag matrix (MA), nucleocapsid (NC), and capsid (CA) domains each play distinct and crucial roles in the formation of the immature virion (Lingappa et al., 2014; Mailler et al., 2016). The MA domain is critical for targeting of Gag to the plasma membrane, and this depends on MA myristoylation and the binding of host RNA. Specifically, binding of transfer RNAs to a basic region in MA is thought to block Gag interaction with internal membranes prior to reaching the highly acidic inner leaflet of the plasma membrane (Gaines et al., 2018). The Arc NTD has predicted homology and several functional similarities to retroviral MA (Campillos et al., 2006). The positively charged Arc NTD confers binding to phospholipid membranes, and Arc binding to plasma membranes is promoted by palmitoylation of a cysteine cluster _94_CLCRC_98_ within the putative endophilin binding site of the second coil (Barylko et al., 2017; Hallin et al., 2018). Here, we found that mutation of the endophilin binding region (s92-98A) containing the palmitoylation sites did not affect Arc oligomerization although it inhibited Arc-mediated transferrin endocytosis in HEK cells. As for studies of Gag assembly at the plasma membrane (Inamdar et al., 2019; Sengupta et al., 2019), super-resolution imaging techniques will be needed to resolve the relationship between Arc attachment to membranes and higher-order oligomerization.

The NC domain has a dual function in binding both viral and host RNA. Recognition of viral genomic RNA sequences requires specific trans-acting zinc-finger motifs (Lingappa et al., 2014; Lu et al., 2011). In contrast, Gag multimerization requires basic residues in the NC. A patch of basic residues at the NC N-terminus mediates non-specific binding to host RNA (Lingappa et al., 2006, 2014). The host RNA is thought to function as a scaffold for Gag-Gag interactions and Gag dimers are considered to be building blocks for Gag lattice assembly (Alfadhli et al., 2005; Burniston et al., 1999; Campbell and Rein, 1999; Campbell and Vogt, 1997; Ma and Vogt, 2004; Muriaux et al., 2002; Zhang et al., 1998). Arc does not have amino acid sequence similarity to the Gag NC. However, using single-molecule imaging we found that GFP RNA acts on purified Arc to mediate nucleation of oligomer formation above the dimer stage, and this response is lost in the second coil mutant. Similar to the NC domain, the Arc oligomerization region (99-126) is flanked by positively charged patches (*SI Appendix, Fig. S2*) which may interact with polyanionic RNA. Thus, the Arc NTD resembles the NC domain in mediating RNA-induced multimerization. It is notable in this regard that replacement of NC with a coiled-coil domain forming a leucine zipper can reinstate Gag multimerization (Crist et al., 2008; Johnson et al., 2002).

Studies of synaptic plasticity in mammalian brain have demonstrated a strict spatial-temporal coupling between Arc RNA and protein expression (Panja et al., 2014; Steward et al., 2015). Arc RNA is found in capsids, and evidence from the Drosophila neuromuscular junction implicates Arc protein binding to Arc RNA in capsid assembly. However, the oligomeric state of Arc involved in RNA interaction is unknown. Our findings strongly suggest that RNA-induced oligomerization of Arc is initiated by the dimer. It will therefore be important to determine whether Arc RNA or other endogenous neuronal RNAs play a role in the early stages of Arc oligomerization and particle assembly. In analogy to Gag, oligomerization and encapsidation of RNA cargo may involve distinct RNA-protein interactions.

The core lattice of the retroviral capsid is formed by dimerization of CA domains, which assemble into a hexameric core (Briggs et al., 2009; Pornillos et al., 2009). Whereas recombinant CA undergoes self-association *in vitro*, this is not the case for the isolated Arc CTD or the CTD lobes (present work and (Hallin et al., 2018)). All regions of Arc lacking the NTD are fully monomeric (Hallin et al., 2018). Moreover, the recombinant rat Arc C-terminal region fails to form regular capsids (Pastuzyn et al., 2018), suggesting that the full-length Arc protein is needed for assembly. In the present study, Arc^s113-119A^ formed a stable dimer and failed to form capsids. The synthetic peptide of the oligomeric region showed a strong oligomerization propensity, forming a hexamer *in vitro*. These findings implicate the Arc NTD in molecular interactions within capsid-like structures. The exact mechanism of dimerization is currently unknown, but this assembly may require the CTD, which has dimerization motifs homologous to retroviral CA. Both SAXS analysis and the single-particle EM reconstruction of the Arc^s113-119A^ variant fit well with the presence of two full-length Arc monomers (Hallin et al. 2018) inside the elongated dimer shape. SAXS analysis of MBP-fused Arc^s113-119A^ further shows N-termini on the outside of the dimer, consistent with CTD involvement at the dimer interface. In the full-length Arc monomer, the NTD and CTD interact, stabilizing the flexible linker region (Hallin et al. 2018). In the dimer, a domain swap could take place, allowing NTD-CTD interactions between monomers (model in *SI Appendix, Fig. S5D*). Taken together, the present work suggests that dimers are basic units for capsid assembly, and these dimers are joined together through additional NTD-NTD interactions.

The diameter of Arc capsids based on DLS and negative-stain EM was 30 nm, with considerable variance. These dimensions agree with studies of recombinant rat and Drosophila Arc capsids (Ashley et al., 2018; Pastuzyn et al., 2018). The number of Arc molecules per capsid, based on DLS analysis of volume and mass, is estimated between 40 to 55 units. This range matches the smTIRF analysis, indicating that most higher-order oligomers contained between 40-70 units with some larger species up to 170 units. A particle size of 60 subunits with a diameter of 30 nm corresponds to the smallest icosahedral structure for viral assembly (Ryu, 2017).

The present results have significant implications for elucidating Arc function. Arc dynamics in synaptic plasticity is consistent with the action of a rapidly degraded monomer or low-order oligomer, while Arc assembled into a capsid shell and destined for release in extracellular vesicles implies a metabolically stable protein. The discovery of Arc capsids raises profound questions regarding mechanisms and subcellular sites of assembly, capture of RNA cargo, and the impact of the released extracellular vesicles at target sites. Our findings provide a molecular basis for dissecting the functions of monomeric and dimeric Arc relative to higher-order assemblies and virus-like capsids. Interestingly, Drosophila Arc proteins (dArc1 and dArc2) have CA-like domains and form capsids, although they lack an N-terminal domain. The critical role of the NTD in mammalian Arc capsid formation indicates significant species differences in the mechanism of capsid formation.

## Materials and methods

### Affinity purification assay

Fluorescently tagged Arc and StrepII tagged Arc were coexpressed in HEK293 cells and purified using Strep-Tactin Sepharose (IBA Lifesciences). Formation of complexes between the indicated Arc constructs was analyzed by immunoblotting following SDS electrophoresis. See SI Appendix for details on cell culture, purification and antibodies.

### Plasmids and protein purification

Full-length wild-type and mutant Arc were subcloned into either the pETZZ_1a vector, holding a His-ZZ tag, or into the pHMGWA vector, resulting in a His-MBP tag, with both having a tobacco etch virus (TEV) protease cleavage site. Proteins were expressed in *E. coli* BL21 cells, and extracted using Ni-NTA. Tags were removed by TEV protease and the untagged proteins were used for analysis by SEC-MALS, DLS, negative-stain EM, and SAXS. For extended description of purification protocols and methods, see SI Appendix.

### Single-molecule TIRF

In brief, Snap Alexa 488 labeled and Avi tagged Arc was biotinylated allowing it to be affixed to a glass cover slide using a PEG-biotin/streptavidin linkage. A region of interest (ROI) was exposed to constant 488 nm excitation, causing molecules to photobleach over time. Discrete photobleaching steps was observed and the number of steps correlated to the size of the oligomers.

For a full description of all materials and methods, see SI appendix.

## ACKNOWLEDGEMENTS

This work was supported by a Research Council of Norway Toppforsk grant (249951) to CRB, grant BFU2016-75984 (AEI/FEDER, UE) from the Spanish Ministry of Economy and Competitiveness to JMV, and grant MEXT, Japan (20240032, 16H02455, 22110006, 18H04733, 18H05434) to Y.H. TIRF imaging was performed in the Light Microscopy Facility and Nikon Center of Excellence at the Institute for Applied Life Sciences, University of Massachusetts Amherst with support from the Massachusetts Life Science Center. We gratefully acknowledge SAXS beamtime and beamline support at Diamond Light Source and SOLEIL.

## Author contributions

M.S.E, O.N and C.R.B conceived the study. M.S.E performed cloning and affinity purification assays, protein purification, transferrin uptake assays, confocal imaging and image analysis. O.N performed cloning and wrote ImageJ macro. S.G, T.H and Y.H. performed FLIM-FRET imaging and analysis. D.L, S.A and T.K performed the crosslinking experiments and analysis. R.O, J.J.C and M.M.S designed and performed the single-molecule imaging experiments. H.J.B, M.I.F and A.M prepared human recombinant Arc for EM imaging and performed DLS analysis. J.C and J.M.V performed EM imaging and 3D reconstruction analysis. E.I.H and P.K performed protein purifications, CD spectroscopy, SEC-MALS, and SAXS experiments. G.M. and R.L performed SAXS on MBP-Arc. M.S.E and C.R.B wrote the paper with contributions from all authors.

## Notes

#### Summary of Updates

Author list updated

