## supplemental figures and tables for "Molecular determinants of Arc oligomerization and formation of virus-like capsids"

### SUPPLEMENTAL INFORMATION APPENDIX

#### Materials and Methods

**Plasmids**. Human wild-type full length Arc was cloned into the pETZZ_1a vector holding a His-ZZ expression tag and TEV cleavage site. The rat Arc sequence with the s113-119A mutant was cloned into the pHMGWA vector, resulting in an N-terminal histidine and maltose binding protein tag fused to the protein when expressed. A TEV protease cleavage site located between the fusion tag and the protein allowed removal of the tag, leaving one serine residue on the N-terminal side of the protein. The Arc constructs used for single-molecule photobleaching experiments were constructed with an N-terminal SUMO-His tag for purification, followed by a Snap tag (Snap26m) for labeling, and an Avi tag at the C-terminal end for biotinylation. Gibson assembly was used to clone mutant Arc 113-119A. The FRET sensors used were described previously (Hallin et al., 2018), all behind the cytomegalovirus promoter. Other plasmids encoding GST- or fluorescent protein-fused Arc fragments were obtained by subcloning corresponding Arc fragments in pGEX-4T-3 (GE Healthcare Life Sciences) or FRET sensor plasmids between BamHI and NotI restriction sites. Plasmids encoding StrepII-tagged Arc fragments were generated from FRET sensors by substituting C-terminal fluorescent protein with StrepII-encoding oligo (GCGGCCGC-A-TCCGGA-TGGAGCCACCCGCAGTTCGAGAAA-GGT-GGA-GGT-TCC-GGA-GGT-GGA-TCG-GGA-GGT-GGA-TCG-TGGAGCCACCCGCAGTTCGAAAAA-GGT-TAA-CTCGAG) between NotI and XhoI restriction sites. Mutagenesis was performed using QuikChange Lightning Multi Site-Directed Mutagenesis Kit (Agilent) according to the manufacturer’s instructions. All constructs were verified by sequencing.

**Protein expression and purification.** All constructs were transformed to BL21 competent cells (Agilent) for IPTG-induced protein expression. *GST-fused Arc and GST.* Cells were lysed in 100 mM NaCl, 10 mM Tris-HCl (pH 8.0), 1 mM EDTA, 1% Triton x-100 (v/v). Cleared lysates were incubated with glutathione-Sepharose beads (GE Healthcare Life Sciences) for 2 hours, beads were washed and bound protein eluted with 10 mM reduced glutathione. Samples were dialyzed overnight against 20 mM

Tris-HCl (pH 7.4), 150 mM NaCl. Purity of proteins were checked by separation on SDS gels and staining with InstantBlue protein stain (Merck) before measuring concentration with a Nanodrop Spectrophotometer (Thermo Fisher Scientific).

*Purification of Arc for smTIRF.* Cells were lysed in Buffer A (25 mM Tris pH 8.5, 150 mM KCl, 1 mM DTT, 50 mM imidazole). Supernatant was filtered and loaded onto a 5 mL Ni-NTA column (all purification reagents were purchased from GE Healthcare Life Sciences). The protein was eluted by using Buffer B (25 mM Tris, 50 mM KCl, 1 mM DTT, 500 mM imidazole). The SUMO-His tag was cleaved by adding Ulp1 protease overnight at 4° and the sample desalted using Buffer C (20 mM Tris pH 7.5, 125 mM NaCl, 2 mM TCEP) on a 26/10 desalting column. The sample was loaded onto a Q-FF 5 mL column and eluted using a gradient of NaCl. The sample was then run over an Superdex 200 size exclusion column in Buffer C.

*Purification of Arc for DLS and negative-stain EM.* Full-length human Arc was expressed in *E. coli* and purified mainly as previously reported (Myrum et al 2015) with modifications (to be published). After TEV cleavage, protein was buffer-exchanged into 20 mM phosphate (pH 7) using a PD10 column.

*Purification of Arc$^{s113\text{-}119A}$ for SEC-MALS, DLS, SAXS and EM.* Cells were harvested, washed with a solution of 100 mM Tris-HCl (pH 7.5) and 170 mM NaCl, lysed in a buffer containing 40 mM Hepes (pH 7.5), 100 mM NaCl, 1mM DTT, 0.1 g/l lysozyme by one freeze-thaw cycle followed by sonication. The lysed cells were centrifuged at 16 000 g for 30 minutes at 4°C and the supernatant was loaded onto a Ni-NTA resin. The His-tagged protein was eluted using imidazole, treated with recombinant TEV protease and dialyzed against 20 mM HEPES (pH 7.5), 100 mM NaCl, 1 mM DTT for 16 hours at 4°C. The now tagless protein was passed through the Ni-NTA resin again before purification by size exclusion chromatography, using a Superdex S200 16/600 (GE Healthcare Life Sciences) column equilibrated with 20 mM Tris-HCl (pH 7.4) and 150 mM NaCl. The protein of interest gave one homogenous peak which was collected and DTT was added to a final concentration of 1 mM. The protein sample was then concentrated using a 10 kDa MWCO spin concentrator to a concentration of 4.4 g/l. The protein concentration was determined by absorbance measurements at 280 nm.

*Commercial proteins. A* peptide of Arc amino acids 99-132 (Ac-QETIANLERWVKREMHVREVFYRLERWADRLES-OH) and a peptide array of partially overlapping peptides derived from rat Arc (NP_062234.1:71-147) were synthesized by INTAVIS Bioanalytical Instruments.

**Peptide array.** The peptide array was activated in methanol, washed in Tris-buffered saline with Tween (TBST: 50 mM Tris-HCl (pH 7.5), 150 mM NaCl, 0.1% (v/v) Tween 20) and blocked in TBST + 1% BSA (w/v). The array was first incubated with 0.5 μg/mL purified GST for 1 hour at room temperature, washed three times in TBST and electrotransferred to nitrocellulose membrane using the Trans Blot Turbo transfer system (Bio-Rad Laboratories). After transfer the array was stripped in regeneration buffer (62.5 mM Tris-HCl (pH 6.7) + 2% SDS (w/v)) for 30 min, washed three times in TBST and blocked before the second round of incubation with 0.5 μg/mL GST-fused Arc 78-140 followed by electrotransfer. Transferred proteins were detected by an anti-GST antibody by Western blotting using Pierce ECL Western Blotting Substrate (Thermo Fisher Scientific) according to the manufacturer's instructions.

**Size-exclusion chromatography – multi-angle light scattering (SEC-MALS).** The absolute molecular mass of the Arc oligomerization region peptide 99-132 and $Arc^{s113\text{-}119A}$ mutant were determined with SEC-MALS, using a miniDAWN Treos MALS detector. The SEC columns used were a Superdex S75 Increase 10/300 for the peptide and a Superdex S200 Increase 10/300 for the $Arc^{s113\text{-}119A}$ mutant. The running buffer consisted of 20 mM Tris-HCl (pH 7.6) and 150 mM NaCl. The SEC-MALS system was calibrated with ribonuclease A for the peptide experiment and bovine serum albumin for the $Arc^{s113\text{-}119A}$ mutant. The protein concentration was measured with an online refractometer.

**Circular dichroism spectroscopy.** The ellipticity of Arc peptide 99-132 was recorded using a Jasco J-810 Spectropolarimeter (JASCO Products Company) and a 1-mm quartz cuvette. The protein concentration was 0.2 g/l in a buffer consisting of 20 mM phosphate (pH 7.6). The measurements were done at +20°C.

**Dynamic light scattering.** The hydrodynamic diameter of $Arc^{WT}$ and $Arc^{s113\text{-}119A}$ were measured using dynamic light scattering (DLS) at 1 mg/ml at 4 °C in the 20 mM phosphate (pH 7) and 20 mM Tris-HCl (pH 7.6) and 150 mM NaCl, respectively. A Malvern Zetasizer Nano ZS with a HeNe laser at 633 nm was used with a fixed scattering angle of 173°. The Malvern DTS software was used to evaluate the intensity- and volume size distributions, and estimate kDa.

**Small-angle X-ray scattering.** SAXS data collection for the purified recombinant non-tagged Arc$^{s113-119A}$ protein was done on the B21 beamline at Diamond (Oxfordshire, UK) and on the SWING beamline at SOLEIL (Gif-sur-Yvette, France) for the MBP-tagged s113-119A mutant protein. The data were collected using a SEC-SAXS setup, where SAXS frames are collected as the protein elutes from a SEC column. The columns used were Shodex KW404-4F at Diamond and an Agilent ProSEC-300S at SOLEIL. The running buffer used was 20 mM Tris-HCl (pH 7.4) with 150 mM NaCl. The protein concentration was 8 mg/ml for the untagged protein and 20 mg/ml for the MBP-tagged protein. SAXS data for the oligomerization domain were collected at SOLEIL in batch mode at a concentration of 13 mg/ml in a buffer consisting of 20 mM Tris-HCl (pH 7.4) with 150 mM NaCl. All SAXS measurements were done at +10°C. SAXS data were processed using ATSAS (Franke et al. 2017), and the collected frames were checked to avoid radiation damage. SAXS models were generated using DAMMIN (Svergun, 1999), DAMMIF (Franke and Svergun, 2009), and GASBOR (Svergun et al., 2001).

**Single-molecule TIRF (total internal reflection fluorescence).**

*Coverslip preparation.* The Attofluor cell chamber (ThermoFisher) and coverslips were washed and sonicated in 1% Hellmanex, and then washed and sonicated again in 50% isopropanol. The coverslips and donuts were then air dried. On the day of the experiment, the coverslips were plasma cleaned and then secured into the donuts. 250 μL of PEG-Biotin and PEG-PLL (1mg/mL; 500: 3 ratio of the two solutions by volume) were added to the coverslip and incubated for 30 min. The coverslips were washed in 1x PBS. All liquid was removed from the coverslip and 250 μL of streptavidin (10 mg/mL) was added to coverslips and incubated for 30 min. The coverslips were then washed in 20 mM Tris (pH 7.5), 125 mM NaCl, 2 mM TCEP and incubated for another 10 min followed by wash with Buffer C.

*Sample preparation.* Purified Arc was labeled with SNAP Alexa 488 dye (ThermoFisher). The dye was added to the protein in 2 molar excess and allowed to react in the dark for 1 hour at RT. The sample was then desalted into Buffer C using a PD25 column. For experiments without RNA addition, Alexa488-labeled Arc was diluted to 3 nM in Buffer C and added onto the prepared coverslip. After a 1 min incubation, the coverslip was washed 7x with Buffer C and an oxygen scavenger was added to the coverslip (5% glucose, 0.5 mg/mL glucose oxidase, 40 mg/mL catalase). For experiments with RNA, 5 μM Arc was mixed with 0.1 mg/ml EGFP mRNA in RNAse free water for 2 h. The samples were then diluted and added to the prepared coverslip as above. For the Alexa dye control, a biotin-conjugated 488 dye (Biotium #80019) was dissolved in 25 mM Tris (pH 8.5) and diluted to 3 nM with PBS and added to prepared coverslips as above.

*Single-molecule TIRF microscopy.* The sample was imaged on the NSTORM/TIRF microscope housed in the UMass IALS Light Microscopy Facility. A total of 10 randomly chosen 1048x1048 pixel fields of view (164 x 164 μm) were imaged on each coverslip using the 488 nM laser at 30% power with an exposure time of 10 ms. Using the Nikon 100X Plan Apo TIRF objective and a Hamamatsu sCMOS camera, images were recorded for 15 seconds and no pixels were saturated. *Photobleaching analysis.* TIRF images were analyzed using the NIS-Elements (Nikon, version 5.02) software package. First, each 15 second time-lapse series was cropped to include the first 5 seconds in order to attain maximum intensity projections. From these images, single particles (ROIs) were selected based on threshold, shape, and size using the General Analysis module. For the analysis of lower-order oligomers, a maximum intensity projection image in time was created over the first 5 seconds. The minimum threshold was 131 AU. Higher-order oligomers were selected by using the initial intensity of the first frame with a minimum intensity of 285. Intensities for each single particle were calculated over the trajectory of the 15 seconds photobleaching time-lapse. Fluorescence intensity vs. time plots were manually analyzed to 1) calculate representative step size and 2) discard traces with evidence of fluorophore blinking or irregular patterns such as single frame events or ROIs that had no initial intensity above baseline. The initial intensity for each ROI was calculated from averaging the first 50 time points. The baseline fluorescence was calculated by averaging the last 100 data points. Finally, the number of photobleaching steps was calculated using the following equation: (initial intensity-baseline intensity)/step size. Plots were generated using Prism 7.

**Negative-stain electron microscopy (EM) and 3D single-particle reconstruction.**

*Sample preparation.* Purified Arc samples were either directly incubated on the EM grids, or subjected to SEC on a Superdex 200 10/300 GL column (GE Healthcare Life Sciences) equilibrated with PBS (pH 7.4). For the latter, 250 μl fractions of the main peak (600-200 kDa) were collected and analyzed by negative-staining EM. Where *GFP mRNA* is indicated, 2 mg/ml Arc samples were incubated with 7.3% RNA (as in Pastuzyn et al., 2018; CleanCap EGFP mRNA (Trilink BioTechnologies)) for 30 min, diluted to a suitable concentration for EM and subsequently incubated on the EM grids. In all cases, carbon-coated 200-mesh copper/rhodium grids were glow discharged for 15 sec in a vacuum chamber at 15mA. 5 μL samples were applied to the grids and incubated for 1 min, and excess sample was blotted with Whatmann paper. Grids were washed with phosphate buffer once, then stained with 2% uranyl acetate and blotted again to remove the excess contrast agent.

*Image acquisition and processing*. For $Arc^{WT}$, images were taken using a Tecnai $G2^2$ FEG 200 (FEI) microscope operated at 200 kV and equipped with a 4k x 4k FEI Eagle CCD camera at a nominal magnification of 50,000X. In the case of the Arc 113-119A mutant, images were taken using a JEOL 1010 JEM electron microscope operated at 80kV and equipped with a CCD camera (4Kx4K TemCam-F416, TVIPS). Images were recorded at a 65,000X nominal magnification with a pixel size of 15.50 μm (2.4 Å/px sampling rate). These images were processed following the Scipion processing workflow (de la Rosa-Trevín et al., 2016). Images were CTF-corrected using CTFFIND4 (Rohou and Grigorieff, 2015). A total of 55,057 particles were automatically selected using Xmipp and 2D-classified using Relion 2.0 (Kimanius et al., 2016) and CL2D (Sorzano et al., 2010). Some of the best classes were used as a template to build an initial model using RANSAC (Vargas et al., 2014). This model was filtered to 60 Å and used for refinement with Relion 3D auto-refine of the 28871 particles selected from the 2D classification, rendering a 21 Å model. The apparent C2 symmetry of this first model was confirmed by applying this symmetry in the auto-refine process, after which a 3D reconstruction with a similar shape and resolution was generated. Visualization of the 3D models and docking of the atomic structures into EM volumes was performed manually using USCF Chimera (Pettersen et al., 2004).

**Cell culture, hippocampal slice-culture preparation and transfection.** HEK293FT cells (R70007, Thermo Fisher Scientific) were grown in Dulbecco's modified Eagle's medium supplemented with 10% FBS and penicillin/streptomycin (Sigma-Aldrich) at 37 °C in a humidified incubator with 5% $CO_2$. For imaging studies, cells were plated on poly-L-lysine coated coverslips. Cells were transfected using Lipofectamine™ 2000 Transfection Reagent (Thermo Fisher Scientific) according to the manufacturer’s instructions. Transverse hippocampal slice cultures were prepared from Sprague Dawley rats as described before at P8-10 and maintained for 7-10 days before transfection (Hallin et al., 2018). Ballistic DNA transfection was performed using a Gene Gun (Helios); 1.6-μm gold microcarriers (Bio-Rad Laboratories) were coated with plasmid DNA and fired directly into individual wells in 6-well culture dishes containing hippocampal slices.

**Affinity purification assay.** 24 hours after transfection, cells were washed in ice-cold PBS and lysed in buffer containing 25 mM Tris-HCl (pH 7.4), 150 mM NaCl, 1 mM EDTA, 0.5 mM DTT, 0.5% Triton

X-100 (vol/vol), cOmplete Mini Protease Inhibitor Tablets and PhosSTOP Phosphatase Inhibitor Cocktail Tablets (Sigma-Aldrich). After centrifugation, 40 µl of Strep-Tactin Sepharose (IBA Lifesciences) was added to the lysate followed by incubation for 1 hour at 4 °C. Beads were washed in lysis buffer and immobilized proteins eluted by boiling in Laemmli sample buffer prior to separation on polyacrylamide gels by SDS-PAGE and immunoblotting. Probing of membranes with Strep-Tactin®-HRP conjugate was performed according to the manufacturer's instructions.

***In situ* protein crosslinking.** Transfected HEK293 cells were harvested in PBS buffer and incubated for 10 min with the chemical crosslinking reagent Disuccinimidyl glutarate (DSG, Sigma-Aldrich), dissolved in DMSO to a final concentration of 0.1 mM. The reaction was quenched with Tris (pH 7.5) for 10 minutes. Cells were lysed in buffer containing 60 mM Hepes (pH 7.5), 150 mM NaCl, 1 mM EDTA, 1mM DTT and cOmplete Mini Protease Inhibitor for 20 minutes at 4℃. Protein concentrations of cleared lysates were determined with the Pierce™ BCA Protein Assay Kit (Thermo Fisher Scientific) and the same amounts of protein were loaded prior to SDS-PAGE and immunoblotting. The intensity of bands was measured with ImageJ. One-way ANOVA with Bonferroni multiple comparison was used for statistical analysis.

**Antibodies.** Primary antibodies; mouse anti-GFP Antibody (B-2) (Santa Cruz Biotechnology), Strep-Tactin®-HRP conjugate (IBA Lifesciences), mouse anti-Arc (Santa Cruz Biotechnology), goat anti-GST (GE Healthcare Life Sciences). Secondary antibodies; HRP-conjugated anti-mouse and anti-goat antibodies from Merck.

**Transferrin uptake assay and image analysis.** HEK293FT cells plated on poly-L-lysine coated coverslips were serum starved for 4 hours and placed on ice for 10 min prior to incubation with 15 µg/ml Transferrin-AF647 (Thermo Fisher Scientific) for 15 min at 37 °C. Cells were washed and subsequently fixed in 4% PFA before mounting in ProLongGold with DAPI (Thermo Fisher Scientific). Images were acquired on a Leica SP5 confocal microscope with a 40x objective. Settings were kept constant for each sample. Acquired stacks were analyzed by a custom-written macro in ImageJ. In brief, channels were split and used to create binary masks for transfected and non-transfected cells. Individual transfected cells were segmented using the watershed algorithm, and their masks were created using "Analyze Particles" function. All images were manually checked after the automated analysis to exclude overlay masks for overlapping transfected and non-transfected cells. Intensity measurements for every channel were then performed for every transfected cell. Statistical significance between samples was tested by Kruskal-Wallis with Dunn's multiple comparisons post-hoc correction after normalization tests. The ImageJ macro is available upon request.

**FLIM-FRET imaging.** Imaging and data analysis were performed as described previously (Hallin et al, 2018), with the same positive and negative controls for FRET. Briefly, two-photon FLIM-FRET imaging was performed on an Olympus FV1000 with an SPC-830 (Becker & Hickl) photon counting board for time-correlated single-photon counting. Photons were counted for 10–20 s, depending on expression level, in 64 × 64 pixels. The donor (EGFP) was excited using a 910-nm 2-photon laser (Spectra-Physics), and emission was captured in an H7422-40 detector (Hamamatsu). Fluorescence lifetime was calculated in SPC-image software (Becker & Hickl), using mono-exponential curve fitting of photon distributions.

**Secondary structure predictions.**

*In silico* secondary structure predictions were performed using the COILS (Lupas et al., 1991) and JPred (Drozdetskiy et al., 2015) servers.

#### SUPPLEMENTAL TABLES AND FIGURE LEGENDS

**Table S1.** Deletion mutants (red) and substitution mutants (blue) in the Arc NTD second coil.

**Fig. S1. Interaction between Arc second coil (78-140) regions.**

StrepII-tagged Arc 78-140 was coexpressed with mTq2-fused full-length Arc or residues 78-140. StrepII-tagged Arc interacted with full-length and second coil Arc. Due to the small size of StrepII-tagged 78-140 we were not able to detect it by western blot analysis. Cell lysates were incubated with Strep-Tactin Sepharose and the bound proteins eluted in sample buffer. Proteins were detected after SDS-PAGE and Western blot analysis by anti-GFP and anti-Strep antibodies

**Fig. S2. Amino acid charges in Arc with oligomerization domain (yellow) and 7-amino acid oligomerization motif (red).**

**Fig. S3. The Arc oligomerization region peptide forms a well-defined hexamer.**

The oligomerization region is amino acid 99-126. In the synthetic peptide (99-132), six C-terminal amino acids were added as definition at the single residue level is uncertain with the methods used (7-ala scanning and peptide tiling arrays).

(A) CD spectrum of Arc peptide 99-132 shows that the peptide adopts an α-helical conformation.

(B) SEC-MALS chromatogram of Arc peptide 99-132 demonstrates the presence of oligomers with molecular mass corresponding to hexamer.

(C) SAXS scattering curve for the Arc peptide. The raw data are shown in blue and the fit of the hybrid model (panel F) as a black line.

(D) Ab initio modeling of various oligomeric chain-like assemblies using GASBOR indicates the presence of ~5.5 Arc peptide monomers in the complex.

(E) Distance distribution analysis shows a maximum dimension of >100 Å for the peptide sample; as a single peptide in a helical conformation (as shown by CD) can span only 40-50 Å, this shows the presence of a 2-fold symmetry axis in the middle, with a head-to-head interaction between peptides.

(F) An Arc peptide hexamer, with strict P32 symmetry, fits the synchrotron SAXS data (panel C) extremely well and concurs with the results from SEC-MALS. A single peptide is shown in blue.

**Fig. S4. Mutation of Arc oligomerization domain blocks transferrin endocytosis in HEK293FT cells**

(A) Representative images of transferrin (TfR-647) uptake assay in HEK293FT cells transfected with mTq2-Arc. Nuclei stained by DAPI.

(B) Quantification of transferrin uptake in HEK293FT cells represented as mean gray value. Scatter plot with lines indicating median and interquartile range. Kruskal-Wallis test between groups with Dunn's multiple comparisons post-hoc correction ($p<0,05$) shows significant differences compared to wild-type. N=3, n=125, 132, 206, 190, 172, 229, respectively.

**Fig. S5 SAXS analysis of Arc s113-119A mutant protein with and without N-terminal MBP tag.**

(A) SAXS scattering profiles for purified recombinant rat Arc s113-119A mutant protein, MBP-tagged and non-tagged, with fits of generated *ab initio* dummy atom models in red.

(B) SAXS distance distribution functions for purified recombinant rat Arc s113-119A mutant protein, MBP-tagged (dashed) and non-tagged (solid).

(C) Dummy atom models of purified recombinant rat Arc s113-119A mutant protein, MBP-tagged (gray) and non-tagged (green), and purified recombinant human Arc wild-type monomeric protein (yellow). Below the yellow DAMMIN model is a CORAL hybrid model of purified recombinant human Arc wild-type monomeric protein. Yellow models come from reference (Hallin et al., 2018). Bottom right: the SAXS-based average *ab initio* models of the Arc dimer with and without the MBP tag indicate the position of the MBP moiety in the dimer, and hence, the position of the Arc N-terminus. The result shows that the NTD is on the outside of the Arc dimer.

(D) A domain-swapped model for the dimer fits the currently available data. The predicted location of the oligomerization region is indicated by a red bar.

### Supplemental Tables and Figures

**Table S1. Arc sequence depicting deletion mutants (red) and alanine substitution mutants (blue) of the NTD second coil.**

| 10 | 20 | 30 | 40 | 50 | 60 |
|---|---|---|---|---|---|
| MELDHMTTGG | LHAYPAPRGG | PAAKPNVILQ | IGKCRAEMLE | HVRRTHRHLL | TEVSKQVERE |
| 70 | 80 | 90 | 100 | 110 | 120 |
| LKGLHRSVGK | LENNLDGYVP | TGDSQRWKKS | IKACLCRCQE | TIANLERWVK | REMHVWREVF |

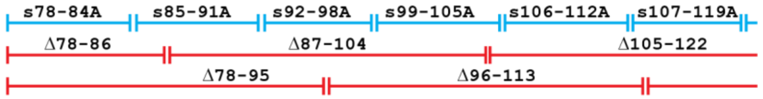


| 130 | 140 | 150 | 160 | 170 | 180 |
|---|---|---|---|---|---|
| YRLERWADRL | ESMGGKYPVG | SEPARHTVSV | GVGGPEPYCQ | EADGYDYTVS | PYAITPPPAA |

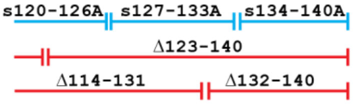


| 190 | 200 | 210 | 220 | 230 | 240 |
|---|---|---|---|---|---|
| GELPEQESVG | AQQYQSWVPG | EDGQPSPGVD | TQIFEDPREF | LSHLEEYLRQ | VGGSEEYWLS |
| 250 | 260 | 270 | 280 | 290 | 300 |
| QIQNHMNGPA | KKWWEFKQGS | VKNWVEFKKE | FLQYSEGTLS | REAIQRELDL | PQKQGEPLDQ |
| 310 | 320 | 330 | 340 | 350 | 360 |
| FLWRKRDLYQ | TLYVDAEEEE | IIQYVVGTLQ | PKFKRFLRHP | LPKTLEQLIQ | RGMEVQDGLE |
| 370 | 380 | 390 | | | |
| QAAEPSVTPL | PTEDETEALT | PALTSESVAS | DRTQPE | | |

**Figure S1. Interaction between Arc second coil peptides (78-140).**

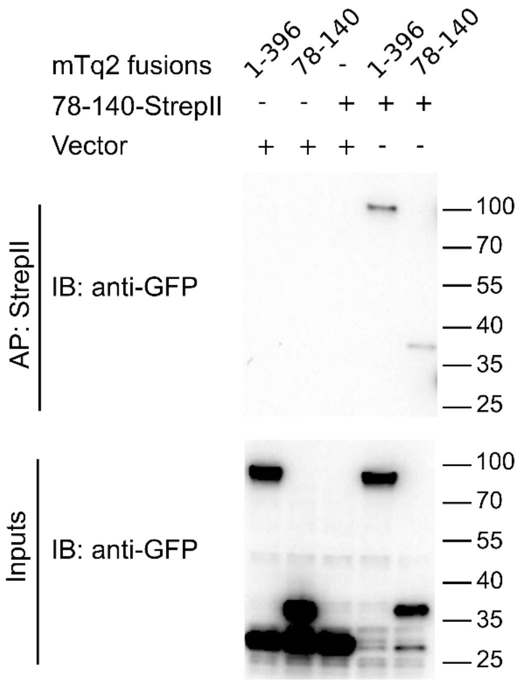


**Figure S2. Isoelectric point of amino acid residues in rat Arc.**
Yellow=oligomerization region (aa 99-126).
Red = oligomerization motif (aa 113-119).

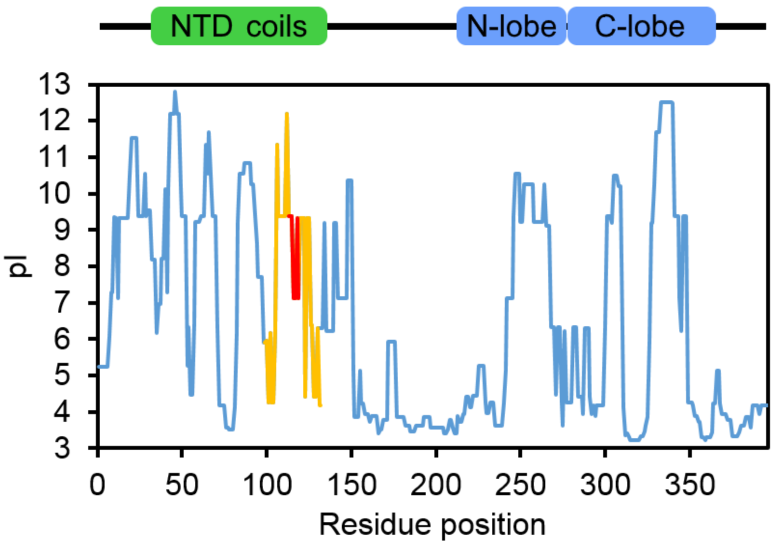

### Figure S3. The oligomerization region peptide forms hexamers

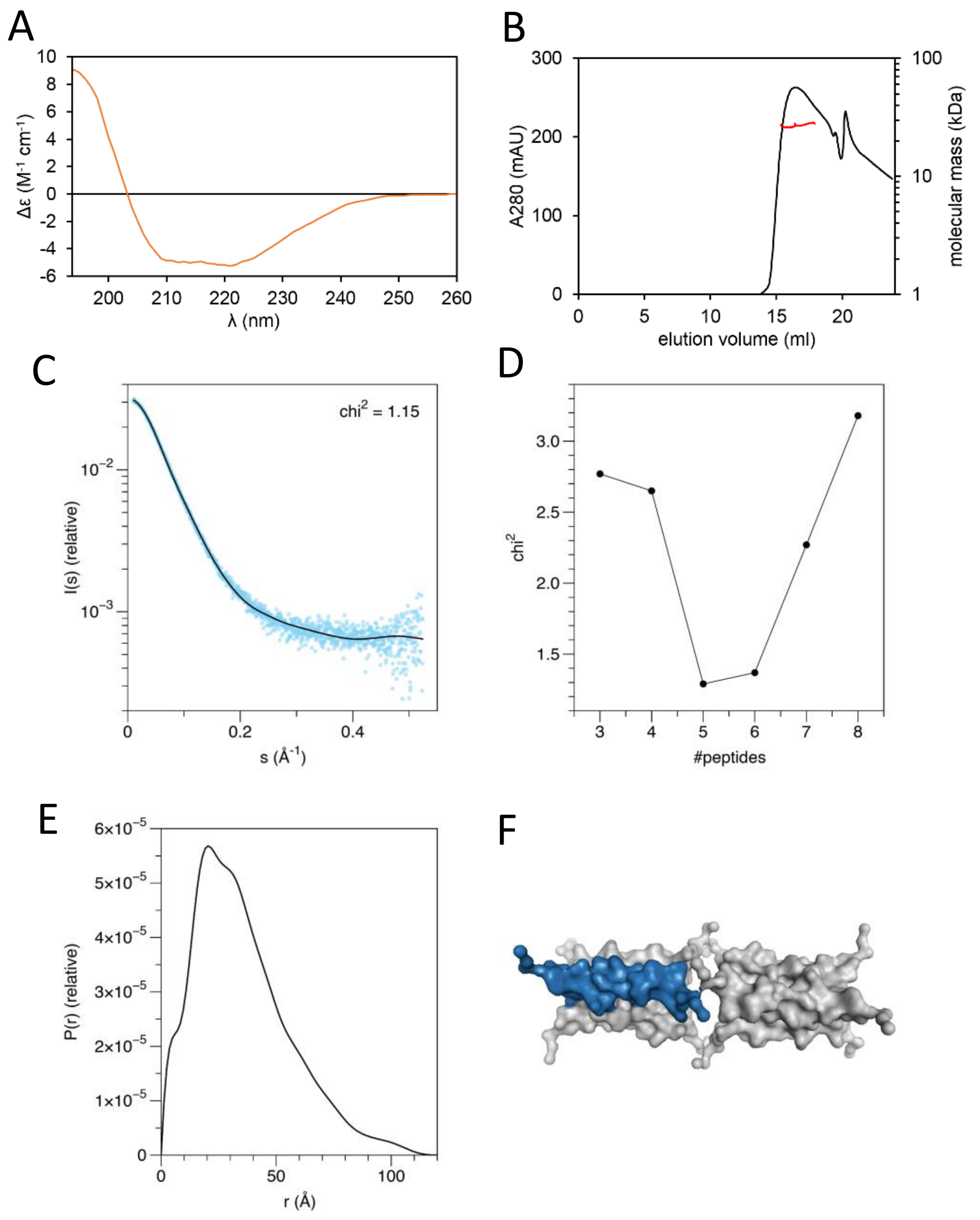

**Figure S4. Mutation of oligomerization region inhibits transferrin endocytosis.**

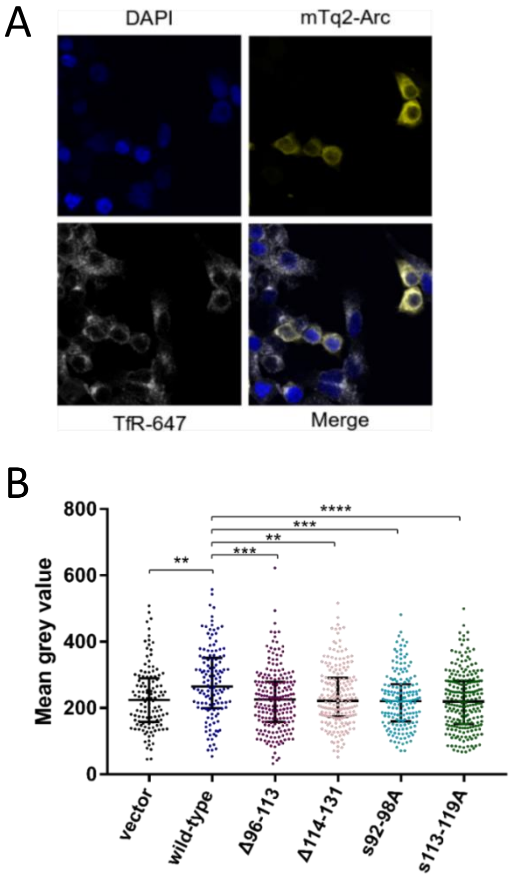

### Figure S5.  SAXS analysis of Arc s113-119A

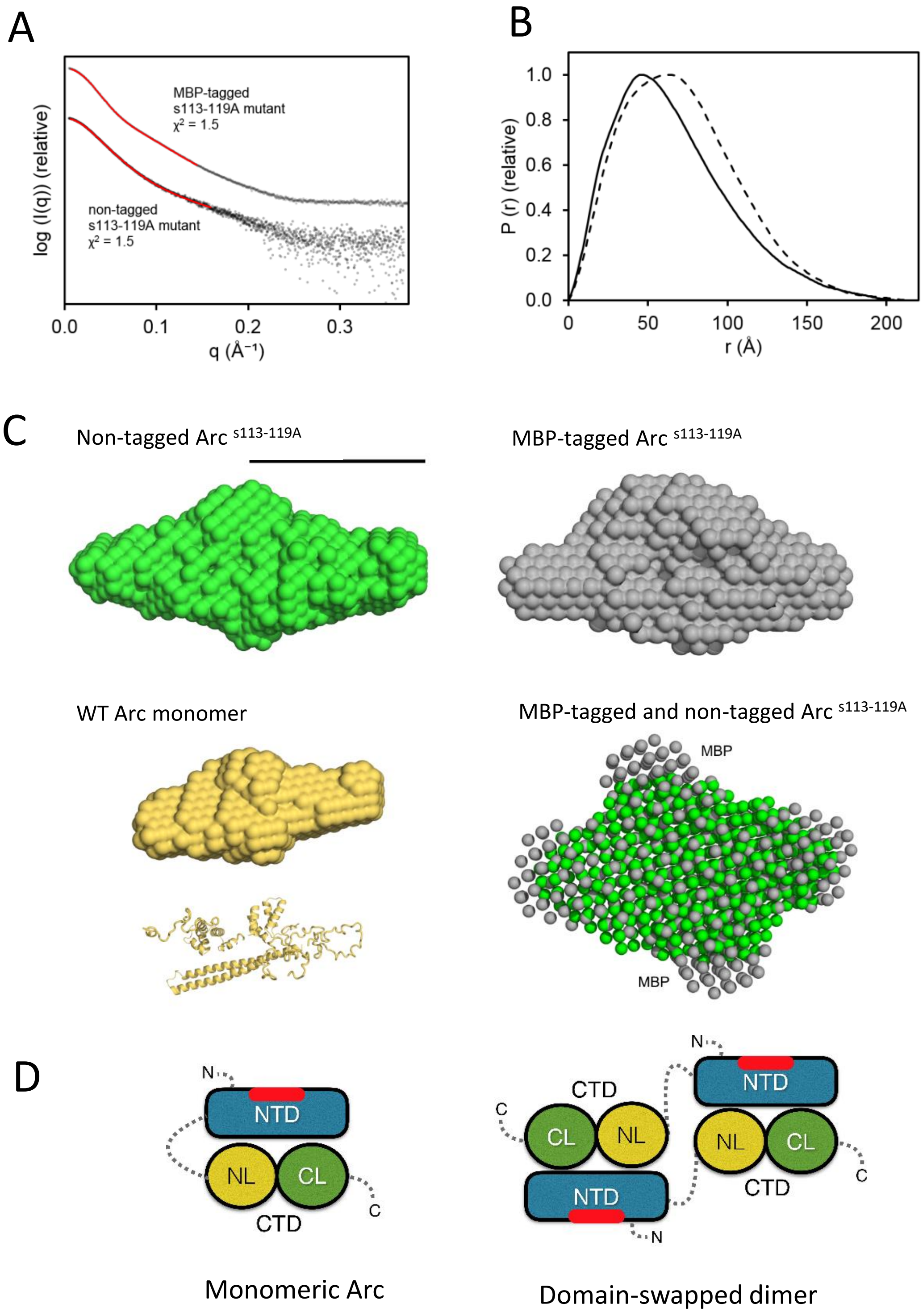
